# Multi-ancestry genome-wide gene-sleep interactions identify novel loci for blood pressure

**DOI:** 10.1101/2020.05.29.123505

**Authors:** Heming Wang, Raymond Noordam, Brian E Cade, Karen Schwander, Thomas W Winkler, Jiwon Lee, Yun Ju Sung, Amy R. Bentley, Alisa K Manning, Hugues Aschard, Tuomas O Kilpeläinen, Marjan Ilkov, Michael R Brown, Andrea R Horimoto, Melissa Richard, Traci M Bartz, Dina Vojinovic, Elise Lim, Jovia L Nierenberg, Yongmei Liu, Kumaraswamynaidu Chitrala, Tuomo Rankinen, Solomon K Musani, Nora Franceschini, Rainer Rauramaa, Maris Alver, Phyllis Zee, Sarah E Harris, Peter J van der Most, Ilja M Nolte, Patricia B Munroe, Nicholette D Palmer, Brigitte Kühnel, Stefan Weiss, Wanqing Wen, Kelly A Hall, Leo-Pekka Lyytikäinen, Jeff O’Connell, Gudny Eiriksdottir, Lenore J Launer, Paul S de Vries, Dan E Arking, Han Chen, Eric Boerwinkle, Jose E Krieger, Pamela J Schreiner, Stephen S Sidney, James M Shikany, Kenneth Rice, Yii-Der Ida Chen, Sina A Gharib, Joshua C Bis, Annemarie I Luik, M Arfan Ikram, André G Uitterlinden, Najaf Amin, Hanfei Xu, Daniel Levy, Jiang He, Kurt K Lohman, Alan B Zonderman, Treva K Rice, Mario Sims, Gregory Wilson, Tamar Sofer, Stephen S Rich, Walter Palmas, Jie Yao, Xiuqing Guo, Jerome I Rotter, Nienke R Biermasz, Dennis O Mook-Kanamori, Lisa W Martin, Ana Barac, Robert B Wallace, Daniel Gottlieb, Pirjo Komulainen, Sami Heikkinen, Reedik Mägi, Lili Milani, Andres Metspalu, John M Starr, Yuri Milaneschi, RJ Waken, Chuan Gao, Melanie Waldenberger, Annette Peters, Konstantin Strauch, Thomas Meitinger, Till Roenneberg, Uwe Völker, Marcus Dörr, Xiao-Ou Shu, Sutapa Mukherjee, David R Hillman, Mika Kähönen, Lynne E Wagenknecht, Christian Gieger, Hans J Grabe, Wei Zheng, Lyle J Palmer, Terho Lehtimäki, Vilmundur Gudnason, Alanna C Morrison, Alexandre C Pereira, Myriam Fornage, Bruce M Psaty, Cornelia M van Duijn, Ching-Ti Liu, Tanika N Kelly, Michele K Evans, Claude Bouchard, Ervin R Fox, Charles Kooperberg, Xiaofeng Zhu, Timo A Lakka, Tõnu Esko, Kari E North, Ian J Deary, Harold Snieder, Brenda WJH Penninx, James Gauderman, Dabeeru C Rao, Susan Redline, Diana van Heemst

## Abstract

Long and short sleep duration are associated with elevated blood pressure (BP), possibly through effects on molecular pathways that influence neuroendocrine and vascular systems. To gain new insights into the genetic basis of sleep-related BP variation, we performed genome-wide gene by short or long sleep duration interaction analyses on four BP traits (systolic BP, diastolic BP, mean arterial pressure, and pulse pressure) across five ancestry groups using 1 degree of freedom (1df) interaction and 2df joint tests. Primary multi-ancestry analyses in 62,969 individuals in stage 1 identified 3 novel loci that were replicated in an additional 59,296 individuals in stage 2, including rs7955964 (*FIGNL2/ANKRD33*) showing significant 1df interactions with long sleep duration and rs73493041 (*SNORA26/C9orf170*) and rs10406644 (*KCTD15/LSM14A*) showing significant 1df interactions with short sleep duration (P_int_ < 5×10^−8^). Secondary ancestry-specific two-stage analyses and combined stage 1 and 2 analyses additionally identified 23 novel loci that need external replication, including 3 and 5 loci showing significant 1df interactions with long and short sleep duration, respectively (P_int_ < 5×10^−8^). Multiple genes mapped to our 26 novel loci have known functions in sleep-wake regulation, nervous and cardiometabolic systems. We also identified new gene by long sleep interactions near five known BP loci (≤1Mb) including *NME7, FAM208A, MKLN1, CEP164*, and *RGL3/ELAVL3* (P_int_ < 5×10^−8^). This study indicates that sleep and primary mechanisms regulating BP may interact to elevate BP level, suggesting novel insights into sleep-related BP regulation.

## Introduction

Hypertension (HTN), including elevations in systolic blood pressure (SBP) and/or diastolic blood pressure (DBP), is a major risk factor for cardiovascular diseases, stroke, renal failure and heart failure ^1,2^. The heritability of HTN is estimated to be 30-60% in family studies ^3,4^. Recent well-powered large genome-wide association studies (GWAS) of blood pressure (BP) have identified over 1,000 loci; however, in total these explain less than 3.5% of BP variation ^5-17^. Gene-environment (G×E) interaction analyses have been proposed to explain additional heritability and identified novel loci for traits associated with cardiometabolic diseases^18-21^.

Long and short sleep durations are associated with elevated BP, possibly through effects on molecular pathways that influence neuroendocrine and vascular systems ^22,23^. Recent multi-ancestry interaction analyses between genetic variants and sleep duration (gene-sleep for short) on blood lipid traits have identified novel loci and potentially distinct mechanisms for short- and long-sleep associated dyslipidemia, and suggest a modification effect of sleep-wake exposures on lipid biology ^21^. We hypothesize that differences in sleep duration may also modify the effect of genetic factors on BP. Assessment of genetic association studies accounting for potential gene-sleep interactions may help identify novel BP loci and reveal new biological mechanisms that can be explored for treatment or prevention of HTN.

Within the Cohorts for Heart and Aging Research in Genomic Epidemiology (CHARGE) Gene-Lifestyle Interactions Working Group ^24^, we investigate gene-sleep interactions on BP traits in 122,265 individuals from five ancestry groups. We utilize both the 1 degree of freedom (df) test of interaction effect and the 2df joint test of main and interaction effects ^25^, and identify novel loci for BP.

## Results

### Genome-wide gene-sleep interaction analyses

We performed genome-wide meta-analysis of gene-sleep interactions on four BP traits (SBP, DBP, mean arterial pressure [MAP], and pulse pressure [PP]) in 30 cohorts in two stages (Supplementary Notes). Stage 1 discovery analyses included 62,969 individuals of European (EUR), African (AFR), Asian (ASN), Hispanic (HIS), and Brazilian (BRZ) ancestries from 16 studies (Supplementary Tables 1-3). Stage 2 replication analyses included 59,296 individuals of EUR, AFR, ASN and HIS ancestries from 14 additional studies (Supplementary Tables 4-6). We examined long total sleep time (LTST) and short total sleep time (STST) separately as lifestyle exposures. Given the heterogeneity of age, sleep duration and BP levels across cohorts and ancestry groups, as well as differences in how sleep duration was assessed (Supplementary Tables 2 and 5), we followed procedures used in prior research ^21^ to categorize 20% of each sample as long sleepers and 20% as short sleepers based on responses to questionnaires, accounting for age and sex variability within each cohort (Methods).

**Table 1.**
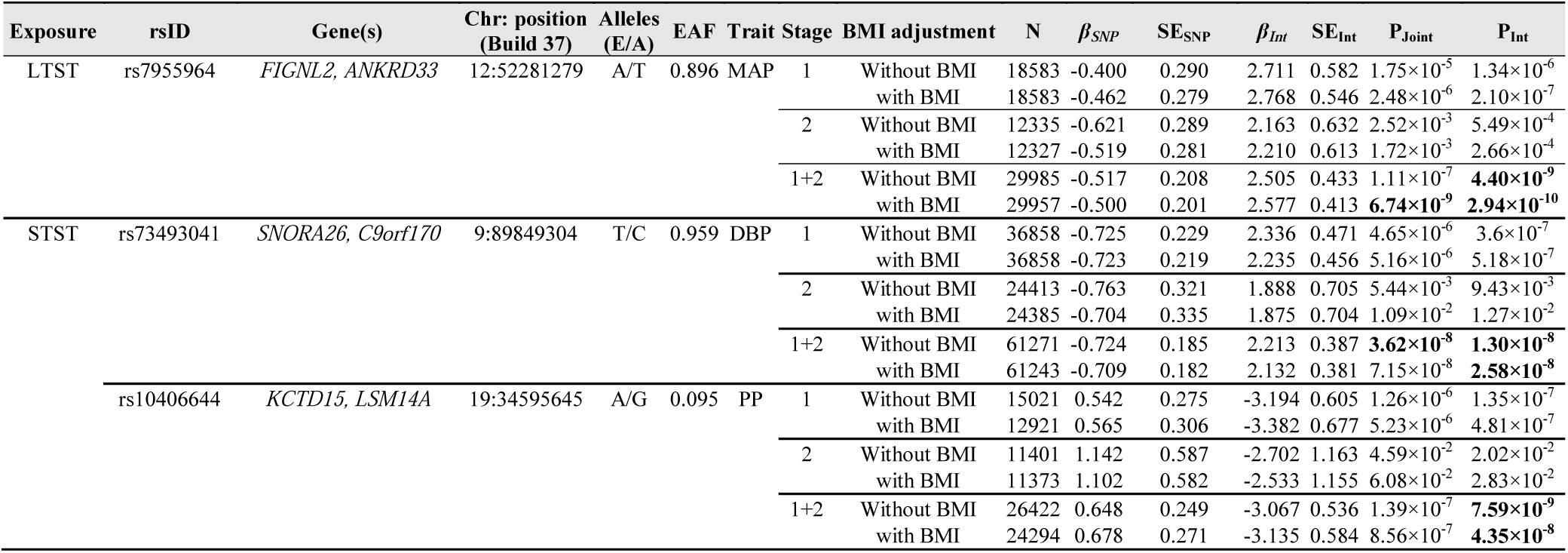
Replicated novel BP loci identified incorporating gene-sleep interactions with and without BMI adjustment.

The overall study design of primary multi-ancestry analyses and secondary analyses restricted to EUR and AFR groups is described in Fig. 1. The analyses were performed without and with adjustment for body mass index (BMI) to identify genetic loci through or independent to obesity pathways (Methods) ^26,27^. The Miami and QQ plots of stage 1 discovery analyses are provided in Supplementary Figs 1-6. Variants with P_int_ or P_joint_ <10^−6^ in stage 1 (n_variant_=1976) were followed up in stage 2 replication analyses and subsequently meta-analyzed with stage 1 summary statistics. Of these, 1081 variants were available in stage 2 cohorts and passed quality control, of which 268 variants showed P_joint_ or P_int_ <0.05. The replication significance threshold was defined as stage 2 P<0.05 and stage 1 + 2 P<5×10^−8^, with consistent directions of association effects.

**Fig. 1.**
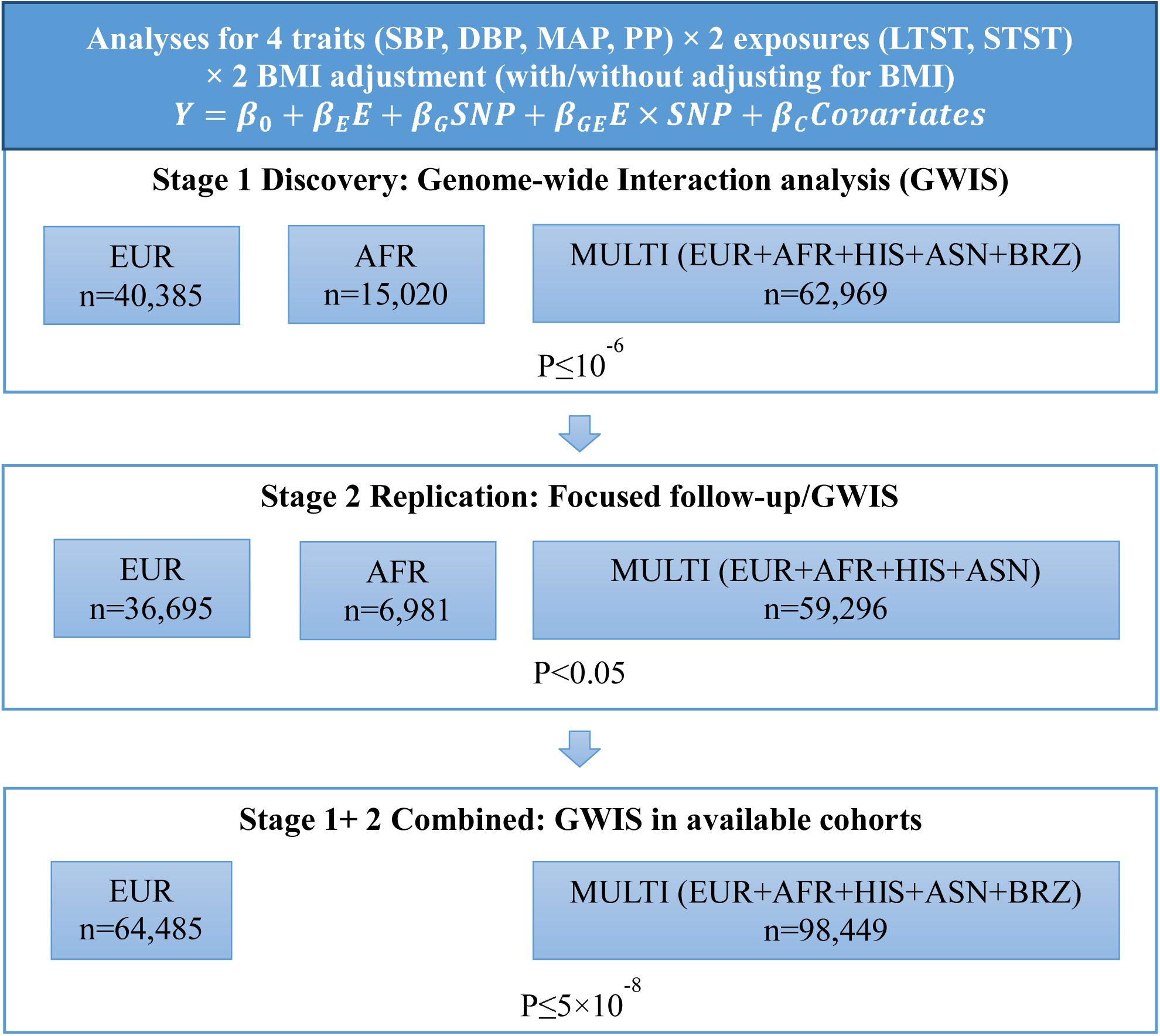
Study overview.

Our primary two-stage multi-ancestry analyses replicated 3 novel loci (r^2^<0.1 and >1Mb from any previously identified BP locus) using the 1df interaction test. Among these loci, rs7955964 (*FIGNL2/ANKRD33*) showed significant 1df interactions with LTST, while rs73493041 (*SNORA26/C9orf170*) and rs10406644 (*KCTD15/LSM14A*) revealed significant 1df interactions with STST (P_int_<5×10^−8^; Table 1). The regional association plots are shown in Supplementary Fig. 7. Consistent directions of main and interaction effects were observed across different cohorts (Fig. 2). Note that since rs7955964 and rs10406644 are common only in AFR and HIS cohorts (minor allele frequency > 1%), the association effects may be driven by African ancestral alleles. With and without adjustment for BMI showed similar association results at those three loci (Table 1). Using 2df joint test we identified 9 loci that were within 1Mb from previously reported BP loci (P_joint_<5×10^−8^; Supplementary Table 7). However, none of these loci showed nominal significant 1df interaction association with LTST or STST (P_int_>0.05).

**Fig. 2.**
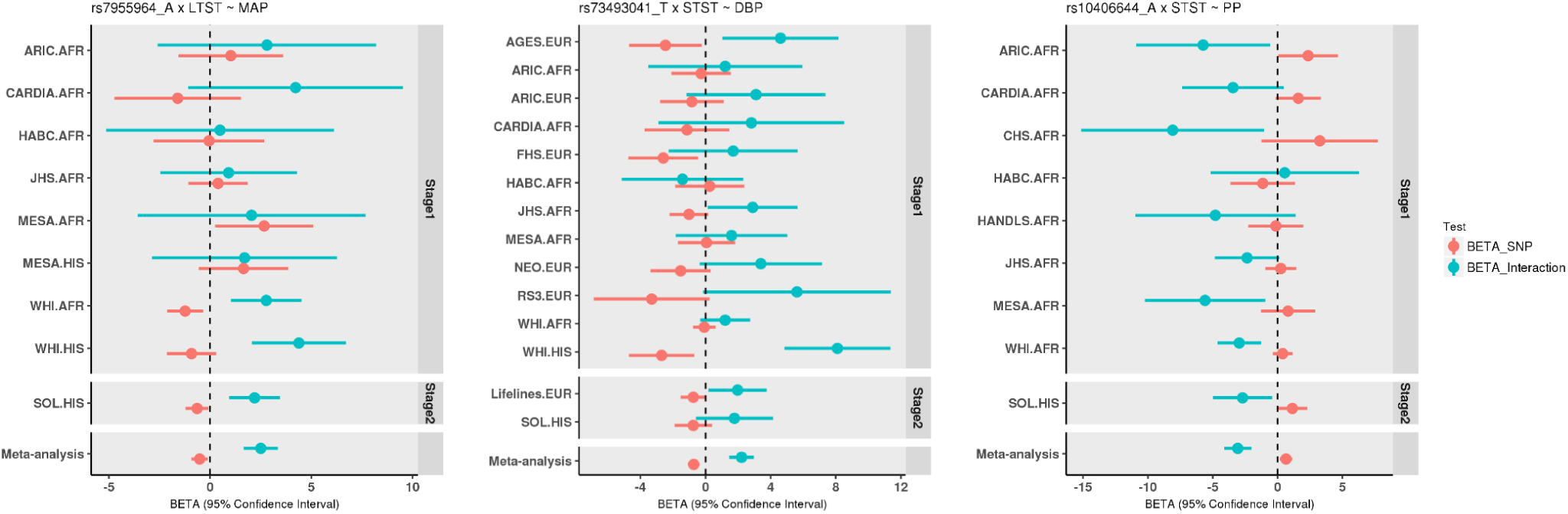
Forest plots of 3 replicated novel loci identified in genome-wide gene-sleep 1df interaction analyses in multi-ancestry population.

In secondary meta-analyses restricted to EUR or AFR individuals, the 1df interaction test did not identify any significant locus. The 2df joint test in EUR group identified 1 locus near previously reported BP loci (≤1Mb; Supplementary Table 7). The 2df joint test in AFR group identified 3 novel loci after incorporating interactions with LTST (stage 1 P_joint_ <5×10^−8^ and stage 2 P_joint_<0.05, with consistent directions of main effects), including rs111887471 (*TRPC3/KIAA1109*), rs114043188 (*ANK*), and rs138967672 (*RP11-322L20*.*1/RP11-736P16*.*1*) (Supplementary Table 8 and Supplementary Fig. 8). However, these 3 variants did not survive our formal replication criteria of stage 1+2 P<5×10^−8^, possibly reflecting heterogeneity between discovery and replication cohorts.

We further performed multi-ancestry genome-wide meta-analyses combining stage 1 and stage 2 cohorts to maximize statistical power (Methods; Miami and QQ plots in Supplementary Figs 9-12). These analyses additionally identified 20 loci near previously reported BP loci (≤1Mb) and 56 unreported loci (>1Mb from previously reported BP) with P_joint_ or P_int_<5×10^−8^ (Supplementary Tables 9-11). Loci near known BP genes, including *NME7, FAM208A, MKLN1, CEP164*, and *RGL3/ELAVL3* ^8-12^ showed significant 1df interactions with LTST (P_int_<5×10^−8^; Supplementary Table 9). Replication in independent datasets is needed to validate those unreported loci. We only report 20 novel loci that passed a stricter significance threshold (P_joint_ or P_int_ <3.125×10^−9^), accounting for 2 independent BP traits, 2 exposures, 2 tests (joint and interaction), with and without BMI adjustment (Supplementary Table 10 and Supplementary Fig 13). Among these 20 novel loci, 3 loci showed genome-wide significant 1df interaction with LTST and 5 loci showed genome-wide significant 1df interactions with STST (P_int_<5×10^−8^; Supplementary Table 10). We also looked up the previously validated 362 BP loci ^5-16^ and 113 sleep duration loci ^28^ in the combined analyses, but none of these showed significant 1df interactions after accounting for multiple comparisons (P_int_>10^−4^; Supplementary Tables 12-15).

We estimated the variance of BP traits explained by 3 replicated and 23 additional novel loci (3 in AFR two-stage analyses and 20 in combined analyses) using the R package VarExp ^29^ (Supplementary Table 16). The 3 replicated novel variants together explained 0.002-0.039% and 0.304-0.939% of BP variation in EUR and AFR, respectively. The other 23 variants additionally explained 0.230-0.419% and 0.806-2.597% of BP variation in EUR and AFR. Given the limited sample sizes in the AFR group, the estimation of BP variation in AFR is likely inflated.

### Associations with other relevant traits

We looked up the associations between the 3 replicated and 23 additional novel loci with cardiovascular diseases, stroke, chronic kidney disease, and self-reported and objective (derived from 7-day accelerometry) sleep traits using publicly available genome-wide summary statistics from large GWAS ^28,30-39^ (Supplementary Tables 17-22). One of the replicated loci rs73493041 (*SNORA26/C9orf170*) was associated with self-reported chronotype (morningness vs eveningness) (P=9.1×10^−6^; Supplementary Table 21). Among the other novel loci, rs17036094 (*PSRC1/MYBPHL*) was associated with coronary artery disease and myocardial infarction (P≤0.005; Supplementary Table 18), and rs140526840 (*FSTL5*) was associated with chronic kidney disease (P=0.006; Supplementary Table 20),

### Bioinformatics analyses

All of the 26 novel variants were mapped to intergenic or intronic regions using HaploReg ^40^, including 4 in promoter histone marks, 11 in enhancer histone marks, 10 in DNAse, 3 altered the binding sites of regulatory proteins and 2 conserved elements (Supplementary Table 23).

Among the 3 replicated novel loci, rs73493041 (*SNORA26/C9orf170*) was an eQTL for *GAS1* in suprapubic skin using GTEx (v8) ^41^ (Supplementary Table 24). Using PLINK pruning and SNPsea ^42^, rs7955964 (*SNORA26/C9orf170*) was mapped to a region of 10 genes (Supplementary Table 25), including *ANKRD33* and *NR4A1*, implicated in sleep-wake control regulation and the neurovascular system ^43,44^. Rs10406644 (*KCTD15/LSM14A*) was mapped to a region overlapping with 9 genes (Supplementary Table 26), including *KCTD15* and *CHST8*, previously associated with adiposity traits and involved in neurodevelopmental and neuropsychiatric diseases ^45-47^ (see Discussion).

Among the other 23 novel loci, 4 variants showed strong eQTL evidence across various tissues such as blood and adipose tissue (Supplementary Table 24). 14 loci were mapped to genes with known functions in cardiac and nervous systems (e.g., *TRPC3* ^*48*^, *RYR2* ^*49*^, *ANK2* ^*50*^, *GJA4* ^*51*^ and *SORT1*^*52*^) and associated with other cardiometabolic (e.g., *HTR1A* ^53^, *PSRC1* ^54^, *PSKH1* ^55^), inflammatory (e.g. *IL33* ^56^), cognition (e.g., *FRMD4A* ^57^) and psychiatric traits (e.g. *NFATC3* ^58^) (Supplementary Tables 25 and 26).

In total, 11 novel loci harbored genes implicated in Mendelian syndromes such as ventricular tachycardia and cryptogenic cirrhosis. 13 loci harbored one or more genes with potential drug targets (Supplementary Tables 25 and 26).

We performed tissue and pathway enrichment analyses using annotated genes under novel association regions using FUMA ^59^ (Supplementary Tables 27 and 28). Genes under the association regions in gene-LTST interaction analyses were enriched in multiple artery and cardiac muscle related pathways (Supplementary Table 29).

## Discussion

We performed gene-sleep interaction analyses on BP using 122,265 individuals from 5 ancestry groups in 30 studies, using both a 1df test of interaction effect and a 2df joint test of main and interaction effects. Following a two-stage design, we identified 3 novel loci that were replicated in additional samples, including rs7955964 (*FIGNL2/ANKRD33*) showing significant interactions with LTST, and rs73493041 (*SNORA26/C9orf170*) and rs10406644 (*KCTD15/LSM14A*) showing significant interactions with STST (P_int_<5×10^−8^). Secondary analyses additionally identified 3 novel loci with weak replication evidence in AFR groups using 2df joint test. Combined stage 1 and 2 analyses identified another 20 novel loci after accounting for multiple comparisons (P_joint_ or P_int_<3.125×10^−9^), which require external replication. The associations were largely unchanged after additionally adjusting for BMI. Collectively, these 26 loci explained 0.23-0.43% of BP variation in EUR and 1.33-2.96% BP variation in AFR groups.

The emergence of novel loci after considering gene-sleep interactions suggests an important modifying role of sleep on BP regulation, which involves both central and peripheral regulation (including the brain, adrenal glands, kidneys, and vasculature). Insufficient or short sleep can increase BP through effects on elevating sympathetic nervous system activity and altering hypothalamic-pituitary-adrenal (HPA) axis activities, leading to hormonal changes, endothelial dysfunction, insulin resistance, and systemic inflammation ^22,60^. The mechanisms underlying the association between long sleep duration and BP are less well understood, and may reflect circadian misalignment in a 24-hour period, including disrupted sleep-wake cycle, a misalignment of internal biological clocks with the external environment, and desynchronized central and peripheral clocks in tissues relevant for BP control ^61^. The importance of circadian control of BP is evident by the normal nocturnal decline (“dipping”) in BP. Non-dipping of BP, associated with increased mortality, is observed with both sleep disturbances and abnormalities of sodium transport in the kidney ^62,63^. Our data suggest that sleep and renal and neuro-endocrine control of BP may interact to influence susceptibility to HTN. The novel loci found by gene-LTST and gene-STST interaction analyses were distinct, supporting the different mechanisms of short and long sleep modifying BP. Similarly, in prior gene-sleep interaction analyses for blood lipids, LTST and STST each also modified gene effects in a non-overlapping pattern ^21^.

Primary two-stage multi-ancestry analyses identified 3 novel loci. At rs7955964 (*FIGNL2/ANKRD33*), *ANKRD33* is expressed in retinal photoreceptors and the pineal gland and acts as a transcriptional repressor for CRX-activated photoreceptor gene regulation ^43^. Given the importance of light in the central regulation of circadian rhythms, long sleep- a circadian disruptor-may interact with this gene to influence BP ^62^. Additionally, *NR4A1* (also at rs7955964) is a member of the nuclear hormone receptors, which regulate neurohormonal systems including dopamine and norepinephrine and cardiac stress responses ^44^. At rs10406644 (*KCTD15/LSM14A*), *CHST8* and *KCTD15* are associated with adiposity traits ^45,46^ and *GPI* functions in glucose metabolism and immune system pathways ^64,65^. Short sleep may exacerbate weight gain and metabolic dysfunction ^66^, thus amplifying effects of this locus on BP. *KCTD15* belongs to a gene family involved in neurodevelopmental and neuropsychiatric diseases ^47^. Rs73493041 (*SNORA26/C9orf170*) was an eQTL for *GAS1*, a pleiotropic regulator of cellular homeostasis and widely expressed in the central nervous system ^67,68^. This variant was also significantly associated with self-reported chronotype, an indicator of circadian preference (P=9.1×10^−6^; Supplementary Table 20).

Given the high prevalence of HTN in African Americans, there is a critical need to identify modifiable risk factors. Notably, African Americans have poorly controlled HTN as well as circadian abnormalities in BP regulation ^69^. They also have a higher prevalence of short and long sleep duration compared to individuals of European ancestry ^70,71^, likely due to combinations of social-environmental exposures and genetic and epigenetic susceptibility ^72^. In AFR specific gene-LTST analyses we identified 3 unreported loci, including two loci mapped to *TRPC3* and *ANK2* with known functions in cardiac ion (Na^+^ and Ca^2+^) homeostasis ^48,50^. These associations in AFR may reflect differences in BP control with individuals of African ancestry having greater sodium sensitivity^73^, with BP effects amplified by disrupted circadian rhythm regulation due to long sleep ^63^. While intriguing, these associations were observed in African Americans (P_joint_<5×10^−8^) and the only available replication sample was individuals of African ancestry from the UK (P_joint_<0.05). There may be a systematic difference of the admixture structure between the discovery and replication cohorts. Future analyses using larger AFR samples will be needed to validate these findings and further identify the potential for sleep disturbances to interact with BP regulatory systems in AFR who are at increased risk for HTN-associated morbidities.

Although we did not observe significant 1df interactions with sleep duration on the 362 previously reported BP or 101 sleep duration variants (Supplementary Tables 12-15), we identified 30 loci within 1Mb from previously reported BP regions (Supplementary Tables 7 and 9). Among those, variants in/near *NME7, FAM208A, MKLN1, CEP164*, and *RGL3/ELAVL3* showed significant 1df interactions with LTST (P_int_<5×10^−8^; Supplementary Table 9). Some of these genes have known functions in neuronal systems, including *MKLN1* regulating the internalization and transport of the GABA_A_ receptor ^74,75^ and *ELAVL3* encoding a neural-specific RNA-binding protein involved in neuronal differentiation and maintenance ^76^.

In this study we defined short and long sleep duration using self-reported questionnaires, which can result in misclassification ^77,78^, potentially reducing statistical power. Although we used a within cohort approach for harmonizing sleep duration that accounted for age and sex differences across cohorts, there may be systematic residual differences in sleep assessments that resulted in heterogeneity across our samples. Future work using objective measurements (e.g., polysomnography and actigraphy data) may provide further insight into sleep-related BP mechanisms.

Some of our most interesting findings - and ones with high potential public health impact due to the burden of extreme sleep duration and HTN in AFR group. Unfortunately, limited samples of AFR were available for replication. We identified 1,976 variants with significant association effect in gene-sleep interaction analyses in stage 1. However, only 1,081 of those variants were available in stage 2 analyses. Most of the unavailable variants in stage 2 had been identified in non-EUR cohorts and were rare in EUR populations (MAF<1%). Future studies following-up these “missing” variants in diverse groups and additional studies of minority populations are needed to further understand mechanisms for BP regulation that are modulated by sleep. In addition, some of our findings were mapped to large genomic regions covering many genes. Further fine-mapping analyses using sequencing data or biochemistry experiments may further clarify the causal variants.

In summary, we performed a large-scale gene-sleep interaction meta-analyses in multi-ancestry groups. In addition to identifying interactions at a genome-wide significant level near 5 previously reported BP regions, we identified 3 novel loci with formal replication and 23 novel loci that need external replication. Multiple genes showing significant interactions with long or short sleep duration were functional in tissues relevant for BP control, and associated with circadian, metabolic, and neuropsychiatric traits. Prior research indicates that extreme sleep durations (short or long) are strongly associated with cardiovascular disease and mortality through causal and noncausal pathways ^22,23,79^. Our data suggest that interactions between sleep and BP-regulating genes may contribute to the increased cardiovascular morbidity observed with extreme sleep duration.

## Methods

### Samples

This study included 30 cohorts comprising of five ancestry groups including European (EUR), African (AFR), Asian (ASN), Hispanic (HIS), and Brazilian (BRZ) groups. Ancestries were harmonized by each cohort based on self-reported data and removed genetic principal components (PCs) outliers. Gene-sleep interaction analyses were restricted to men and women between 18-80 years old and with available sleep, blood pressure (BP), and genotype data imputed to 1000 genome reference panel. Analyses of each ancestry group were performed locally by individual cohorts following a universal analytic protocol. Descriptions of each cohort are available in Supplementary Notes. Sample characteristics are present in Supplementary Table 1-6.

This work was approved by the Institutional Review Board of Washington University in St. Louis and complies with all relevant ethical regulations. For each of the participating cohorts, the appropriate ethics review board approved the data collection and all participants provided informed consent.

### Phenotypes, lifestyle exposures, and covariates

In this study, we focused on two lifestyle variables (short total sleep time [STST] and long total sleep time [LTST]) and investigated their interaction with genetic components on four BP traits (systolic BP [SBP], diastolic BP [DBP], mean artery pressure [MAP], and pulse pressure [PP]).

Resting/sitting SBP and DBP were recorded in mmHg and averaged among multiple readings at the same visit. Effects of anti-hypertensive (BP lowering) medications were addressed by adding 15 mmHg and 10mmHg to SBP and DBP, respectively. We then derived MAP and PP from the medication adjusted SBP and DBP as MAP=DBP + (SBP-DBP)/3 and PP=SBP-DBP. For each of the four BP variables, we winsorized the very extreme BP values to 6 standard deviations (SD)^24^.

Total sleep time (TST) was harmonized from questions similar to “On an average day, how long do you sleep?”. Questions were either asked as “open (free text)” or “multiple choice”. Individuals reported shorter than 3 or longer than 14 hours of TST were further excluded in this study. For most of the cohorts, we performed linear regression on TST adjusting for age and sex in each ancestry group of each cohort and obtained age- and sex-adjusted residuals. Short total sleep time (STST) and long total sleep time (LTST) were then defined as the lowest 20^th^ and highest 80^th^ percentiles of the residual in each ancestry of each cohort. In some cases, cohort-specific definitions (e.g. “multiple choice”) were used to defined STST and LTST.

We adjusted for age, sex, and when appropriate for field center, family and population structures, and other study specific covariates (e.g., sampling weights and census block in HCHS/SOL^80^). Body mass index (BMI; calculated as weight / height^2^ [kilograms/meters^2^]) has been suggested to influence BP and sleep duration ^26,27^, and may play a potential role in gene-sleep interactions on BP. Therefore, we performed additional analyses adjusting for BMI.

### Genotypes

Each study performed genotyping and imputation locally using various platforms and software as described in Supplementary Tables 3 and 6. The cosmopolitan reference panels from the 1000 Genomes Project Phase I Integrated Release Version 3 Haplotypes (2010-11 data freeze, 2012-03-14 haplotypes) was specified for imputation. Variants with minor allele frequency <1% were excluded by each study.

Upon completion of the analyses by each local institution, all summary data were stored centrally for further processing and meta-analyses. We performed study level quality control (QC) on study specific results using the R package EasyQC ^81^ (www.genepi-regensburg.de/easyqc). We performed harmonization of alleles, comparison of allele frequencies with ancestry-appropriate 1000 Genomes reference data, and harmonization of all single nucleotide polymorphism (SNP) IDs to a standardized nomenclature according to chromosome and position. We further excluded SNPs with the product of minor allele count (MAC) in the exposed group (LTST/STST=1) and imputation quality (MAC_1_×imputation quality) less than 20.

### Discovery analysis in stage 1

The discovery analysis includes 40,385 EUR, 15,020 AFR, 2,465 ASN, 4,436 HIS, and 663 BRZ from 16 cohorts (Supplementary Tables 1-3). Each cohort performed genome-wide interaction analyses using the joint effect model

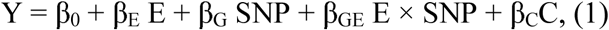

Where Y is one of the four BP phenotypes, E is STST or LTST, G is the dosage of the imputed genetic variant coded additively, and C is the vector of covariates (with or without adjustment of BMI). Therefore, in total 16 analyses in each ancestry group of each cohort were performed. The number of principal components included in the model was chosen according to cohort-specific preferences (ranging from 0 to 10). Various software such as ProbABEL^82^, MMAP and R package sandwich ^83^ were used to estimate the association effects for each study (Supplementary Table 3). Summary statistics of the estimated genetic main effect (β_G_), interaction effect (β_GE_), and a robust estimation of the covariance between β_G_ and β_GE_ were provided by each cohort. We plotted the summary statistics of all effect estimates, standard errors (SE) and p-values across studies to visually compare discrepancies. We also generated SE vs sample and QQ plots to identify analytical problems.

We performed inverse-variance weighted meta-analysis for the 1 degree of freedom (df) interaction term, and 2 df joint fixed-effects meta-analysis of the combined main and interaction effects using Manning et al’s method implemented in the METAL software ^25^. Genomic control correlation was performed before and after meta-analyses. Variants present in less than 3 cohorts or 5000 individuals were further excluded. Ancestry-specific meta-analyses were performed with the same QC criteria in EUR and AFR but not in other groups. We reported results from both 1df interaction (P_int_) and 2df joint effect tests (P_joint_). Variants with P_int_ or P_joint_ <10^−6^ were followed up in phase 2 replication analysis.

### Replication analysis in stage 2 cohort

Stage 2 includes 36,695 EUR, 6,981 AFR, 3,287 ASN, and 12,333 HIS individuals from 14 independent studies (Supplementary Tables 4-6). Seven cohorts performed genome-wide interaction analyses and other cohorts performed focused variant lookup upon the convenience of each analysis team (Supplementary Table 6). Most of the cohorts performed the same joint analysis model (1) as described in the stage 1 analyses, while stratified association analyses of main effect (2) among exposed group (E=1) and unexposed group (E=0) were performed in the UKB AFR individuals.

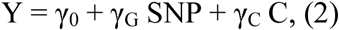

We then estimated the main and interaction effects for the joint regression model as β_G_= γ_G|E=0_ and β_GE_= γ_G|E=1_-γ_G|E=0_ using EasyStrata ^84^. Study level QC was performed similarly as described for stage 1 cohorts. Meta-level QC and genomic control were not performed in stage 2 meta-analyses. Stage 1 and stage 2 summary statistics were also meta-analyzed. The replication threshold was then defined as stage 2 P<0.05 and combined stage 1 + 2 P<5×10^−8^, with similar directions of association effects.

To maximize the statistical power, we also performed genome-wide combined stage 1 and 2 meta-analyses using all stage 1 cohorts and seven stage 2 cohorts (Supplementary Table 6) with genome-wide results in multi-ancestry and EUR groups. Genomic control correlation was performed before and after meta-analyses. Additional novel loci were reported at a stricter significant threshold (P<3.125×10^−9^), accounting for 2 independent BP traits, 2 exposures, 2 tests (joint and interaction), with and without BMI adjustment.

### Bioinformatics analyses

We pruned independently associated loci using the PLINK prune function with r^2^≥0.1 from the lead variant. Loci with physical distance >1MB from any known BP loci were considered as novel. We annotated functional effects for novel loci using HaploReg ^40^, Regulome^85^, and GTex (v8) ^41^ database. In addition to genes mapped by PLINK pruning, we also annotated the genes under the association regions using SNPsea ^42^ with r^2^≥0.5 from the lead variant using 1000 genome European reference panel. Genes under the association regions were further interrogated for associated phenotypes, Mendelian diseases, and druggable targets using PheGeni ^86^, OMIM ^87^, and DGIdb ^88^. Tissue and pathway enrichment analyses were performed using online software FUMA ^59^ MAGMA ^89^ and MsigDB ^90^.

## Acknowledgments

This project was supported by the US National Heart, Lung, and Blood Institute (NHLBI) R01HL118305. H.W. and S.R. was supported by NHLBI R35HL135818. B.E.C. was supported by NHLBI K01HL135405. A.R.B. was supported by the Intramural Research Program of the National Institutes of Health in the Center for Research on Genomics and Global Health (CRGGH). The CRGGH is supported by the National Human Genome Research Institute, the National Institute of Diabetes and Digestive and Kidney Diseases, the Center for Information Technology, and the Office of the Director at the National Institutes of Health (1ZIAHG200362). D.v.H. was supported by the European Commission funded project HUMAN (Health-2013-INNOVATION-1-602757). The CHARGE cohorts was supported in part by NHLBI infrastructure grant HL105756. Study-specific acknowledgments can be found in the Supplementary Notes.

## Author contributions

H.W., B.E.C., and J.L. conducted the centralized data analyses, including quality controls, meta-analyses, and post association lookups and bioinformatics. H.W., R.N., B.E.C., K.S., T.W.W., J.L., Y.J.S., A.R.B., D.C.R., S.R. and D.v.H. were part of the writing group and participated in study design, interpreting the data, and drafting the manuscript. All other co-authors were responsible for cohort-level data collection, cohort-level data analysis and critical reviews of the draft paper. All authors approved the final version of the paper that was submitted to the journal.

